# Automated microarray for single-cell sorting and collection of lymphocytes following HIV reactivation

**DOI:** 10.1101/2023.02.02.526757

**Authors:** Belén Cortés-Llanos, Vaibhav Jain, Alicia Volkheimer, Edward P. Browne, David M. Murdoch, Nancy L. Allbritton

**Affiliations:** Department of Bioengineering, University of Washington, Seattle, Washington, USA; Department of Medicine, Duke University, Durham, North Carolina, USA; Department of Molecular Physiology, Duke University, North Carolina, USA; Department of Medicine, University of North Carolina, Chapel Hill, North Carolina, USA; Department of Microbiology and Immunology, University of North Carolina, Chapel Hill, North Carolina, USA; UNC HIV Cure Center, University of North Carolina, Chapel Hill, North Carolina, USA

**Keywords:** microarrays, time-lapse imaging, single-cell, HIV latency reactivation Abstract

## Abstract

A promising strategy to cure HIV infected individuals is to use latency reversing agents (LRAs) to reactivate latent viruses, followed by host clearance of infected reservoir cells. However, reactivation of latent proviruses within infected cells is heterogeneous and often incomplete. This fact limits strategies to cure HIV which may require complete elimination of viable virus from all cellular reservoirs. For this reason, understanding the mechanism(s) of reactivation of HIV within cellular reservoirs is critical to achieve therapeutic success. Methodologies enabling temporal tracking of single cells as they reactivate followed by sorting and molecular analysis of those cells are urgently needed. To this end, microraft arrays were adapted to image T-lymphocytes expressing mCherry under the control of the HIV long terminal repeat (LTR) promoter, in response to the application of various LRAs (prostratin, iBET151, and SAHA). In response to prostratin, iBET151, and SAHA, 30.5 %, 11.2 %, and 12.1 % percentage of cells respectively, reactivated similar to that observed in other experimental systems. The arrays enabled large numbers of single cells (>25,000) to be imaged over time. mCherry fluorescence quantification identified cell subpopulations with differing reactivation kinetics. Significant heterogeneity was observed at the single cell level between different LRAs in terms of time to reactivation, rate of mCherry fluorescence increase upon reactivation, and peak fluorescence attained. In response to prostratin, subpopulations of T lymphocytes with slow and fast reactivation kinetics were identified. Single T-lymphocytes that were either fast or slow reactivators were sorted, and single-cell RNA-sequencing was performed. Different genes associated with inflammation, immune activation, and cellular and viral transcription factors were found. These results advance our conceptual understanding of HIV reactivation dynamics at the single-cell level toward a cure for HIV.

## 1. Introduction

Human immunodeficiency virus (HIV) remains a significant global health challenge. In 2021, nearly 38 million people were living with HIV, 1.5 million people became newly infected, and 680,000 people died from AIDS-related illnesses.^1^ Tens of millions of people currently require antiretroviral therapy (ART) as there is no cure for HIV. ART does not cure infection, and many anti-retroviral drugs exhibit some toxicity and side effects. A major roadblock towards a HIV cure is the ability of the virus to enter a latent state within host immune cells, thereby forming long-lived viral reservoirs. Latently infected cells can reactivate sporadically after the initial infection, leading to viral rebound if ART is interrupted.^2,3^ Latent HIV largely resides within CD4 T lymphocytes, where it remains transcriptionally silent and invisible to the host immune system.^4^ Another major barrier towards a HIV cure is the rarity of latently infected cells, estimated to be as few as 1 in a million T cells. Given these numerous challenges, current technologies are often limited in their ability to study latently infected cells.^5^ An improved understanding of the kinetics of HIV latency and reactivation is critical for progress towards curing HIV.

Previous studies have focused on the reactivation of latently HIV-infected cells followed immediately by immune clearance tools in a strategy known as “shock and kill” to eliminate viral reservoirs.^2^ Reactivation is achieved through the use of latency reversing agents (LRAs) such as protein kinase C (PKC) agonists (prostratin, bryostatin), mitogen-activated protein kinase (MAPK) agonists (procyanidin trimer C1), inducers of positive transcription elongation factor b (P-TEFb) (JQ1, inhibitor of bromodomain and extra-terminal domain (i-BET) 151), epigenetic modifiers such as histone deacetylase inhibitors (HDACis) (romidepsin, trapoxin, suberanilohydroxamic acid (SAHA)), and others.^6^ Computational studies suggest that a 50% cure rate for people with HIV (PWH) could be achieved if the number of latent cells was reduced by 4 orders of magnitude, a challenging metric.^7^ A second hurdle is the high toxicity of LRAs used under conditions where broad reactivation of HIV occurs.^8^ The development of pharmaceuticals or strategies that selectively target the HIV reactivation process while leaving uninfected cells unscathed will be critical to achieve the elimination of HIV from infected individuals.^2,9^

The study of single cells has accelerated our understanding of cellular physiology in a diverse set of fields such as oncology, neuroscience, virology, and immunology.^10,11^ While assays of pooled cell material have provided many insights into the average behavior of a cell population, there is nevertheless a loss of information since properties of differently behaving cells or cell subpopulations are not identifiable. In particular, rare cells with divergent properties (from the average cell) can have significant impact on an organism yet are completely obscured in bulk measurements. Since HIV reactivation is not uniform across an entire population of latent cells (and does not impact the vast majority of normal cells), approaches to track reactivation within single cells are critical to develop new strategies to understand the dynamics of HIV infection and identify new therapeutics.^12^ To provide insights into HIV reactivation, technologies focusing on protein presence or behavior in single cells such as flow cytometry, mass spectrometry, microdevices, and microscopy, have been used to investigate latency reversing agents in cells, quantify and phenotype the viral reservoir, and to study the importance of T lymphocytes in infected individuals.^13–17^ Of these methods, time-lapse imaging of single cells engineered with fluorescent reporters has been particularly valuable in understanding heterogeneity in reactivation kinetics.^18^ Single-cell RNA-sequencing (RNA-seq), DNA-seq, or assay for transposase-accessible chromatin using sequencing (scATAC-seq) combined with new computational methods and single-cell analysis have also improved the understanding of the mechanisms of HIV latency reversal.^16,19,20^ A key gap in all of the above technologies is the inability to perform time lapse imaging of single reactivating cells within a population followed by single-cell gene expression with linkage of the microscopy-measured reactivation kinetics to that cell’s gene expression profile.

Microraft arrays were developed for imaging of single cells over time followed by isolation of the single cells for downstream assays including gene expression.^21(p)^ In this study, microraft arrays were adapted to track and sort individual lymphocytes with and without applying LRAs. The microraft arrays were optimized to track the latency and kinetics of reactivation in large numbers of single cells, followed by identification of reactivated cells with unique properties or reactivation behaviors. Microraft size and lymphocyte adhesion to the microrafts was optimized for nonadherent lymphocytes. Reactivation was monitored using a T cell line with an integrated mCherry reporter of HIV gene expression (LChIT3.2 cell line). LChIT3.2 cells were reactivated using three different LRAs: prostratin, SAHA (vorinostat), and iBET151. Reactivation kinetics of single cells were tracked over time by microscopy followed by sorting and gene expression analysis of those cells with unique behaviors. The transcriptional heterogeneity of cells that reactivated quickly (fast activators), slowly, or not at all was examined. The optimization and demonstration of the utility of the microraft arrays to HIV-infected cells will advance our conceptual understanding of when and how HIV-infected, single cells exit latency.

## 2. Experimental

### 2.1 Microraft array fabrication

The template for the microraft arrays was formed on a poly(dimethylsiloxane) (PDMS) template using photolithography and soft lithography as previously described.^22^ To form microrafts on the PDMS microwell template, the array was dip-coated into a magnetic polystyrene solution [18.5% mass/mass poly(styrene-co-acrylic acid) in gamma-butyrolactone (GBL, Sigma-Aldrich, B103608) with 1% maghemite iron oxide (γ-Fe_2_O_3_) nanoparticles.^23,24^ The microarray slide was glued to a cassette and detached from its substrate as described previously.^22^ Each microraft array contained 19,600 microwells (100 x 100 x 60 μm, L x W x H; 50 μm spacing between wells).

### 2.2 LChiT3.2 cell line

LChiT3.2 is a human T lymphoblastic leukemia cell line (CEM T cell line) with a single copy of an LTR-mCherry-IRES-Tat cassette.^25^ Cells were maintained in RPMI 1640 (Gibco, 22400-089) with 2 mM L-glutamine and 20 mM HEPES. All media was supplemented with 10% of fetal bovine serum (100-106, GEMINI Bench Mark hiFBS) and 1% penicillin/streptomycin (P/S) (GIBCO, 15140-122), 10,000 units/mL penicillin, 10,000 μg/mL streptomycin.

### 2.3 Cell culture on microraft arrays

Prior to cell culture, the microraft array surface was oxidized in a plasma cleaner (Harrick Plasma, Ithaca, NY) for 5 min to increase surface hydrophilicity. The array was then incubated in ethanol (75% in water) for 30 min. The array was rinsed 3x with phosphate-buffered saline (PBS), 1x for 5 min (Fisher Sci. BF3991). Cell-Tak (354240, Corning) in 0.1 M NaHCO_3_ (144-55-8, Sigma-Aldrich) was adjusted to a pH 6.5-8 using NaOH (1 M) and then added to the microraft array at 3.5 μg protein per cm^2^ of surface area. The microarray with Cell-Tak was incubated at 37 ^0^C for 1 h. The solution was aspirated, and the array was washed 3x with sterile Milli-Q water for 3 min.

Prior to plating cells on the array, cells were stained with the DNA-binding dye, Hoechst 33342, for 30 min at 37 °C (Thermo Fisher). Dead cells were identified by the addition of Sytox Green at 50 nM (Thermo Fisher, S7020). Cells at 1:3 ratio (cells/microrafts) were overlaid onto the microraft arrays in a phenol-red-free medium (RPMI, Thermo Fisher, 11835030, + 2 mM L-Glutamine) supplemented with 10% FBS, 20 mM HEPES and 1% P/S.

### 2.4 Automated imaging

Imaging was performed using a motorized Olympus IX81 inverted microscope (Olympus Corporation, Tokyo, Japan) controlled by custom-built MATLAB software with hardware control using an open-source Micromanager.^24,26^ The microscope was equipped with an MS-2000 motorized stage (ASI, Eugene, OR) and an Orca-Flash 4.0 V2 camera (Hamamatsu, Shizuoka, Japan). The objective used was an Olympus UPLFLN 4x (NA 0.13). Fluorescence imaging of Hoechst 33342, mCherry, and GFP, was performed using DAPI (Chroma ET-DAPI 49000), TxRed (Semrock TxRed-4040B), and FITC (Semrock FITC-3540B) filter sets. Time-lapse images were performed every 2 h for 24 h. The microarray was maintained in focus using a customized software autofocus (MATLAB) employing a modified Laplacian focus algorithm.^21,24^ During time-lapse imaging, a custom-made incubator maintained the cells on the arrays at 37 °C, 60% humidity, and 5% CO_2_.

### 2.5 Image analysis and cell sorting

Individual microrafts were segmented from brightfield images as described previously.^27^ Images were processed using flat field correction and thresholded using Otsu’s method to create a binary mask to segment and identify the microrafts. Each raft was then assigned a unique identifier based on its row and column position. Fluorescence images of the array were collected and processed using a Wiener filter to eliminate noise, top-hat filtering for background subtraction, followed by thresholding using Otsu’s method to identify cells. Individual cells could then be tracked over time in sequentially obtained fluorescence images. A binary mask for each fluorophore (Hoechst, Sytox Green, and mCherry) was created by thresholding, and the fluorescence intensity within the mask borders was calculated for each cell on the array. The area, position, and pixel number were recorded for each cell on a microraft. The fluorescence intensity at each time point was normalized to the mean fluorescence intensity measured at time zero. A k-means algorithm was performed to cluster single cells into groups using fluorescence intensity and time of reactivation.^28,29^ Cells selected for collection were identified as described in the text, and the microraft address for each raft with a selected cell was identified.^27^ A motorized microneedle was mounted on a 4x objective and was moved vertically to pierce the bottom of the PDMS template, dislodging the targeted microraft with the selected cell (attached to the microraft). The microraft with a single cell was then collected using a magnetic wand. The wand was comprised of a magnet (0.8 mm diameter and 10 mm length) mounted within a hollow tube (1 mm diameter, 47 mm length). The wand with the collected microraft/cell was then placed into a PCR tube (RNase free) with a collection buffer (12.5 μl of 10x lysis buffer, RNase inhibitor, 3’SMART-seq CDS Primer IIA, and Nuclease-Free water, SMART-Seq HT kit, Takara, 634455). Once the cell/microraft was collected into a tube, the tube was frozen at −80°C until cDNA synthesis and RNA-seq analysis.

### 2.6 s.c.RNA-seq preparation

Single-cell lysates collected from the microraft array were thawed, and cDNA amplification was performed using the SMART-Seq HT kit (Takara) as directed by the manufacturer. cDNA was quantified using a Qubit (Thermo) and quality validated with a Bioanalyzer (Agilent). Poor quality samples (<1 ng/μl DNA) were excluded from further analysis. The library preparation and sequencing were performed using a Nextera XT Library Prep kit (Illumina) according to the protocol modifications listed in the SMART-Seq HT kit.

### 2.7 s.c.RNA-seq analysis

Fastq files generated after sequencing were first checked for quality using FastQC version 0.11.9. For alignment of sequencing reads to reference transcriptome (GRCh38-2020-A) STAR aligner (version 2.7.2b) was used. Reference transcriptome was first indexed using STAR genomeGenerate function and then sequencing reads for each cell were aligned individually to reference index using STAR alignReads function and genecounts were calculated using quantmode. Resulting gene counts for each cell were concatenated into gene by cell matrix was then imported into R package Seurat (version 4.1.0) for further downstream analysis. Using ScTransform() within Seurat, cells were normalized and top 3,000 genes were selected based on variance using the Seurat implementation FindVariableFeatures(). These genes were then centered and scaled using Seurat’s implementation ScaleData() and percent mitochondrial genes as a variable to regress out. Scaled data was subsequently used for principal components analysis (npcs = 100), and cell cluster analysis was performed using Seurat’s FindNeighbors (dims = 1:5) and FindClusters (resolution = 0.4) functions. Visualization was performed using uniform manifold approximation and projection (RunUMAP, dims = 1:5). Differential gene expression analysis was performed using the Seurat’s FindMarkers function, which uses a Wilcoxon rank sum test and Bonferroni correction to generate adjusted p-values for each gene.

### 2.8 Statistics

All statistical analyses were calculated in GraphPad Prism with * p < 0.05, ** p < 0.01, *** p < 0.001 and ****p<0.0001. Unless otherwise noted, measurements were reported as the mean ± standard deviation of the data points. A two-way ANOVA was used unless specified otherwise. A one-way ANOVA with a Kruskal-Wallis test was performed to compare the slope and end time data for the LRAs. For k-means clusters, the percentage of activated cells was compared by a one-way ANOVA with a Kruskal-Wallis test, while the mCherry fluorescence intensity changes over time between non-activated and activated (fast and slow) cells were compared by a two-way ANOVA with multiple comparisons. Finally, an unpaired t-test was used for comparison of the slope and end point of reactivated fast and slow cells exposed to prostratin.

## 3. Results and discussions

### 3.1 Overview of the platform used to identify and collect cells reactivating in response to LRAs

Microraft arrays were adapted to identify and collect LChIT3.2 T cells based on the kinetics of their HIV LTR-driven mCherry expression in response to LRAs. In this system, the LChIT3.2 cells contain an integrated but transcriptionally silent copy of the HIV LTR promoter, followed by an mCherry fluorescent reporter gene, and the viral Tat transcriptional activator. mCherry expression thus identifies cells undergoing reactivation from HIV latency. For the current application, arrays possessing 19,600 microrafts were fabricated to accommodate up to 6,500 cells per array (Figure 1a,b). Since the LChIT3.2 cells were nonadherent, the arrays were pre-coated with Cell-Tak, a non-immunogenic, adhesive protein, to attach the cells to the surface of the microrafts. Cells on the array were labeled with Hoechst 33342 to mark all cells, and SytoxGreen was added to the media to identify dead cells (Figure 1C). LChiT3.2 cell viability on the arrays was high over the course of the experiments (97.5 ± 1.7%, 8 arrays, 5807 single cells). A small percentage (4.1 ± 2.6, 8 arrays, 5807 single cells) of cells expressed mCherry at 24 h in the absence of an LRA, suggesting that these cells can undergo some spontaneous reactivation.^9,30^ Following addition of an LRA, cells with varying patterns of mCherry expression over time were identified for collection (Figure 1d). The maturation half-life of mCherry is 40 min, a short time with respect to the overall experimental time of 24 h, enabling accurate measurement of viral reactivation.^31,32^ After collecting time-lapse images, the microraft arrays with cells were analyzed using an automated pipeline and cells were clustered into groups based on their mCherry-expression intensity and kinetics. Selected microrafts with cells were released by actuation of a microneedle to move vertically, piercing the PDMS mold and physically dislodging the microraft with its attached cell (Figure 1e). Released microrafts with their cell cargo were collected using a wand with an embedded magnet (Figure 1f). The microraft with its cell cargo was then placed into a collection tube followed by single cell RNA-sequencing (s.c.RNA-seq) (Figure 1g). When microrafts were collected and released, the microraft collection efficiency for this array geometry was 98 ± 3% (3 arrays, 30 microrafts each array). The cell collection efficiency was 67 ± 9% (3 arrays, 30 cells each array) for Cell-Tak adhered cells. Without Cell-Tak-based adhesion, 0 ± 0% (2 arrays, 10 cells each array) of cells could be collected using the microrafts. In prior reports, LaBelle, Attayek and colleagues overlaid gelatin onto the microrafts to encapsulate and trap nonadherent cells within the microraft cup-shaped lumen.^27,33^ Using this method, the cell collection efficiency was 87%; however, this strategy subjects cells to temperature shocks and requires a complex series of media changes at precisely controlled temperatures making the method challenging and time-consuming to automate and execute fast sequential microraft release. For screening and isolating of large numbers of cells (thousands), Cell-Tak-based adhesion was preferable due to its simplicity, automation ease, and low cellular stress.

**FIGURE 1:**
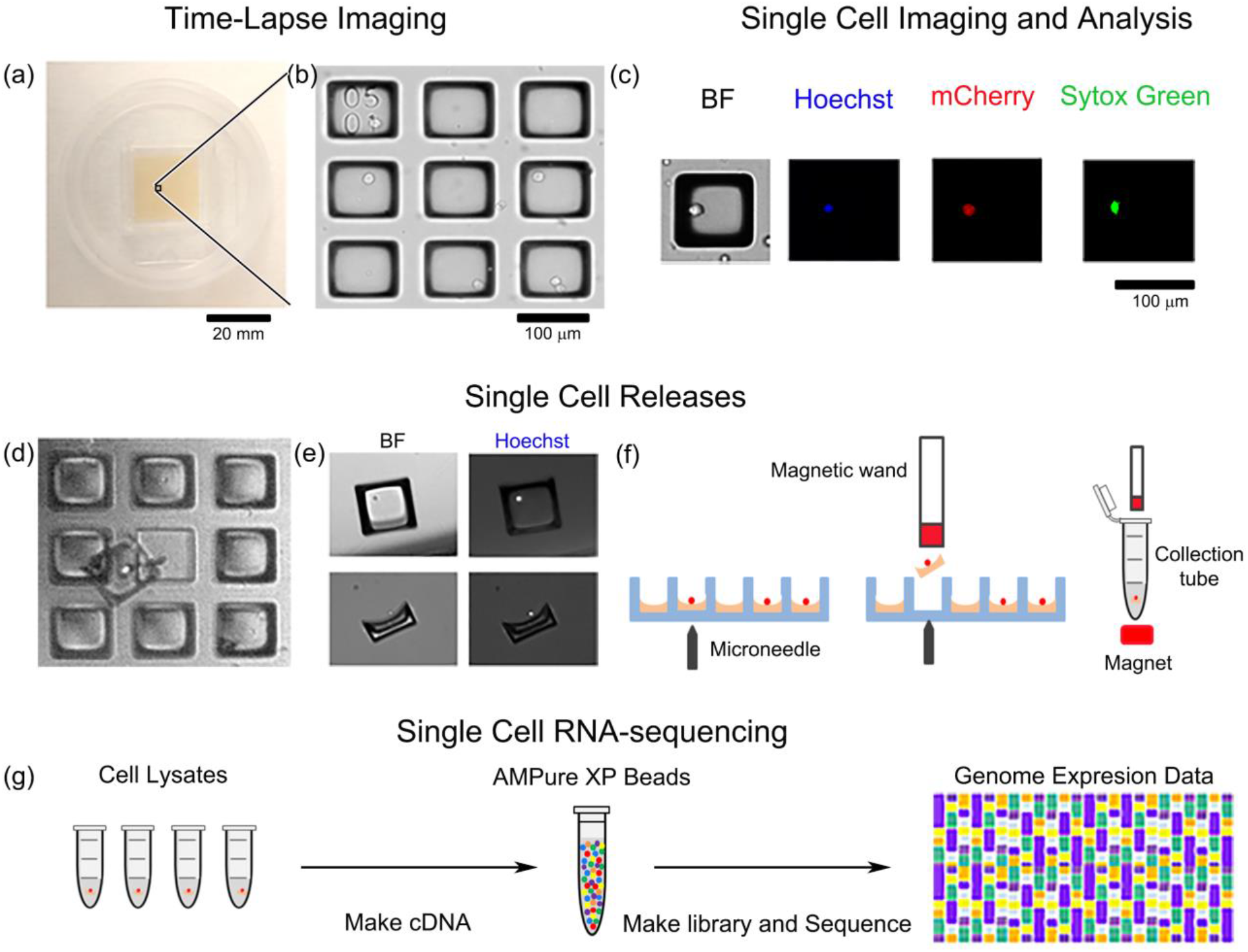
Overview of the automated pipeline to assay gene expression in single cells during drug-induced reactivation of LChIT3.2 cells. (a) Photograph of a microraft array. (b) A section of an array demonstrating 9 microrafts, of which 4 microrafts possessed a single lymphocyte. The upper left corner raft demonstrates a fiducial marker (0501) for array position location. (c) Cells were imaged by brightfield (BF) and fluorescence microscopy to detect mCherry, Hoechst 33342, and Sytox Green fluorescence. Shown are four images of the same raft possessing a single cell. (d) Selected microrafts were released using a microneedle. Shown is a single released microraft that was allowed to settle back down onto the array for imaging. A single cell is attached to the microraft surface. (e) Released microrafts following collection and deposition into a receptacle using a magnetic wand. (f) A schematic of the steps in panels d and e. (g) Collected cells were processed for gene expression analysis.

### 3.2 Cell reactivation using LRAs

LChiT3.2 cells were plated onto a microraft array and imaged over time for expression of mCherry with and without the addition of LRAs. The LRAs prostratin, iBET151, and SAHA each act by a different mechanism. Prostratin is a phorbol ester that activates protein kinase C,^30^ while iBET151 inhibits the family of epigenetic regulators comprised of the bromodomain and extra-terminal (BET) proteins.^34^ SAHA (Vorinostat) acts as a histone deacetylase (HDAC) inhibitor and thus also modulates chromatin configuration.^35^ Cell viability after the addition of the various LRAs was not significantly different from that of the control cells without added LRA (Figure S1). The percentage of cells expressing mCherry fluorescence above a threshold value was significantly greater for cells exposed to an LRA than that of control cells (Figure 2). A significantly greater percentage of cells reactivated in response to prostratin compared to iBET151 and SAHA and the control at 10 h (****p<0.0001) (Figure 2E). By 24 h, the percentage of cells reactivated in response to prostratin was 30.5 ± 5.2%, similar to that observed in studies of human T lymphocytes exposed to prostratin.^9^ These results are also consistent with prior data demonstrating that PKC activators are potent HIV latency reversers.^36^ The percentage of cells reactivated in response to iBET151 and SAHA exposure was significantly greater than that of the control at 18 h (*p=0.0104) and 20 h (**p= 0.0028), respectively. Finally, the peak percentage of cells reactivated at 24 h was significantly lower for iBET51 (11.2 ± 5.3%, ****p<0.0001) and SAHA (12.1 ± 4.3%, ****p <0.0001) relative to the control. The total percentage of cells activated by SAHA was in the range of reactivation percentages observed for SAHA-treated LChit3.2 cells. The percentage of cells activated in response to iBET-151 was similar to that observed for CD4+ T cells infected with latent HIV and exposed to iBET-151. These results suggest that reactivation of the cells on the microraft arrays is similar to that of cells in other latency model systems.^9,37^

**FIGURE 2:**
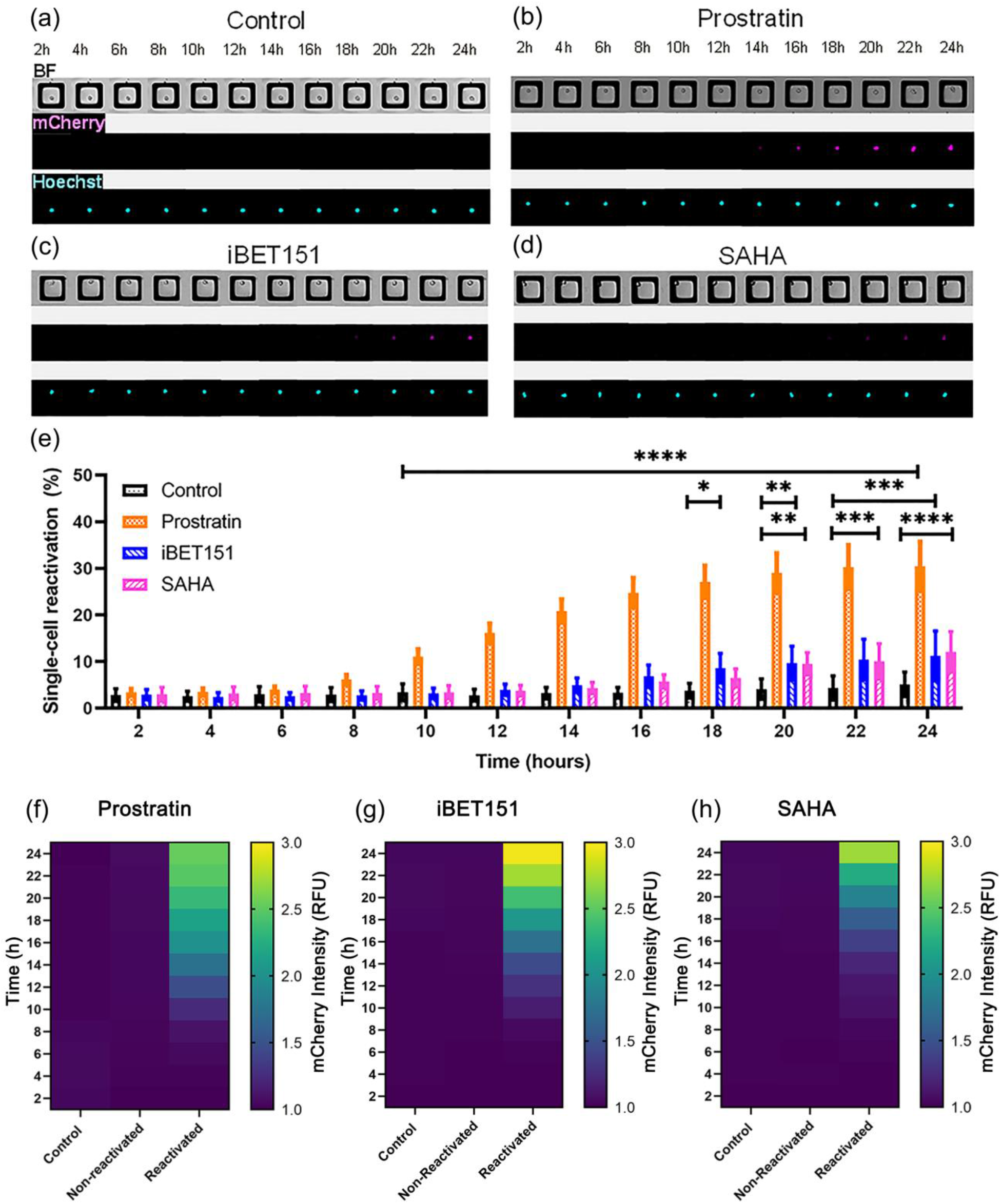
Reactivation of LChiT3.2 cells by LRAs. (a-d) Brightfield and fluorescence images of single cells on the microraft arrays over 24 h without an LRA (a) and with an LRA: prostratin (b), iBET151 (c), and SAHA (d). (e) The total percentage of cells reactivated over time is shown for the unexposed control and LRA-exposed cells. Shown is the average of the data points and the error bars represent one standard deviation (control: n = 8 arrays, with 5807 single cells; prostratin: n = 9 with 8412 single cells; iBET151: n = 4 arrays, 5118 single cells and SAHA: n = 4 arrays, 6354 single cells F-H) Heat maps of mCherry fluorescence intensity for cells exposed to: (f) prostratin-non-reactivated (n=9 arrays, 5657 single cells) and reactivated (n=9 arrays, 2755 single cells), (g) iBET151-non-reactivated (n=4 arrays, 4495 single cells) and reactivated (n=4 arrays, 623 single activated cells) and (h) SAHA-non-reactivated (n=4 arrays, 5902 single cells) and reactivated (n=4 arrays, 452 single activated cells) as well as the control group of cells (unexposed cells, n=8 arrays, 5807 single cells) from each of these experiments.

Based on their peak fluorescence intensity at any time, single cells were divided into two categories: non-activated and activated. Non-activated cells retained their baseline mCherry fluorescence for the entire 24-h of the experiment, while activated cells showed an increase in fluorescence intensity above the threshold during at least one timepoint of the experiment. As expected for each LRA, the fluorescence of the non-activated LRA-exposed cells was similar to that of the control cells never exposed to an LRA (Figure 2f-h). For the activated cells, the mean fluorescence intensity increased over time for each of the LRAs reached a peak at 24 h. On average, the increase in mCherry fluorescence intensity for prostratin-activated cells occurred at shorter experimental times relative to that of the other 2 LRAs, with SAHA-activated cells displaying the longest times to exhibit a mCherry fluorescence increase. For the prostratin-activated cells, a significant increase in the mean mCherry fluorescence was observable by 8 h relative to that of the control cells (**p=0.0012) and non-activated cells (**p=0.0052) (Figure 2f). Using iBET15, activated cells required 10 h compared with the control (***p=0.0007) and non-activated cells (***p=0.0040) to demonstrate a significantly increased mCherry fluorescence (Figure 2g). Finally, cells activated by SAHA exhibited a significant difference in the mean mCherry fluorescence by 12 h compared to the control (*p=0.0424) and 14 h compared to the non-activated cells (***p=0.0003) (Figure 2h). While these averaged data demonstrated the greater speed and potency of prostratin relative to that of iBET151 and SAHA, the data revealed little about the single-cell behavior and importantly how the different cells responded to LRAs.

### 3.3 Single-Cell Heterogeneity During Reactivation

To begin to identify the variation across the single cells in their response to LRAs, the time to reactivation, the time to measurable cellular fluorescence (fluorescence time), and the rate of mCherry increase for each cell was calculated (Table 1). When the behavior of the single cells was averaged together, the fluorescence time for cell reactivation was not statistically different for the 3 LRAs. When examined for each single cell, the range of the fluorescence time displayed in response to each of the LRAs was very large (varying by a factor of 3-8) suggesting that reactivation was governed not only by the LRA but by variable cellular processes or by stochastic behavior of the system. The mCherry fluorescence rate of change once reactivation was initiated was also calculated for each reactivating cell. Prostratin-activated cells, on average increased their rates of fluorescence increase and mCherry fluorescence at a significantly slower rate once activated relative to that for cells activated than iBET151 (Table 1, Figure 3a and e) (*p=0.0338, and p=0.0214, respectively). SAHA activated cells presented a slower mCherry fluorescence rate than iBET151 but it was not significantly different (p=0.1338). But similarly, there was very little variability in the rate of mCherry fluorescence increase once a cell-initiated reactivation. These data suggest that once a cell began to reactivate, each cell reactivated in response to that LRA in a similar manner to other cells exposed to the same LRA. Next, the properties of the cells as they began to activate or at experimental completion was examined for all 3 LRAs. The initial mCherry fluorescence intensity upon activation was similar for cells activating after either short or long fluorescence times, again suggesting that once activated, cells followed similar reactivation pathways in response to a given LRA (Figure 3b-d). Finally, the endpoint mCherry fluorescence intensity was examined for the different LRAs. Prostratin-activated cells possessed a significantly lower final fluorescence intensity than the iBET151-exposed cells (*p=0.0214) but were not significantly different from the SAHA-activated cells (Figure 3E). The higher mCherry increase rate and end point of iBET151-activated cells could be due to the removal of additional blocks to high viral gene expression by this drug. These data demonstrate that activated cells exhibited different behaviors in response to each LRA as they reactivated, possibility related to individual LRA mechanisms of action and activated pathways.

**TABLE 1:** Characteristics of LChiT3.2 cells that were activated in response to an LRA.

| Parameter | Prostratin | iBET151 | SAHA |
| --- | --- | --- | --- |
| **Fluorescence time (h)** |  |  |  |
| Mean (all activated cells) | 15.2 ± 8.9 | 16.5 ± 8.1 | 17.3 ± 7.5 |
| Range observed for single reactivating cells | 4-24 | 6-24 | 8-24 |
| **Rate of mCherry fluorescence increase (RFU/h)** |  |  |  |
| Mean (all activated cells) | 0.13 ± 0.02 | 0.18 ± 0.01 | 0.13 ± 0.01 |
| Range observed for single reactivating cells | 0.1 - 0.15 | 0.17 – 0.19 | 0.13 – 0.14 |
| **Cell reactivation rate (# of cells/h)** |  |  |  |
| Mean (all activated cells) | 15.6 ± 4.7 | 9.6 ± 6.8 | 9.0 ± 7.0 |
| Range observed for single reactivating cells | 10.4 – 24.9 | 3.7 – 17.4 | 3.4 - 16.9 |

**FIGURE 3:**
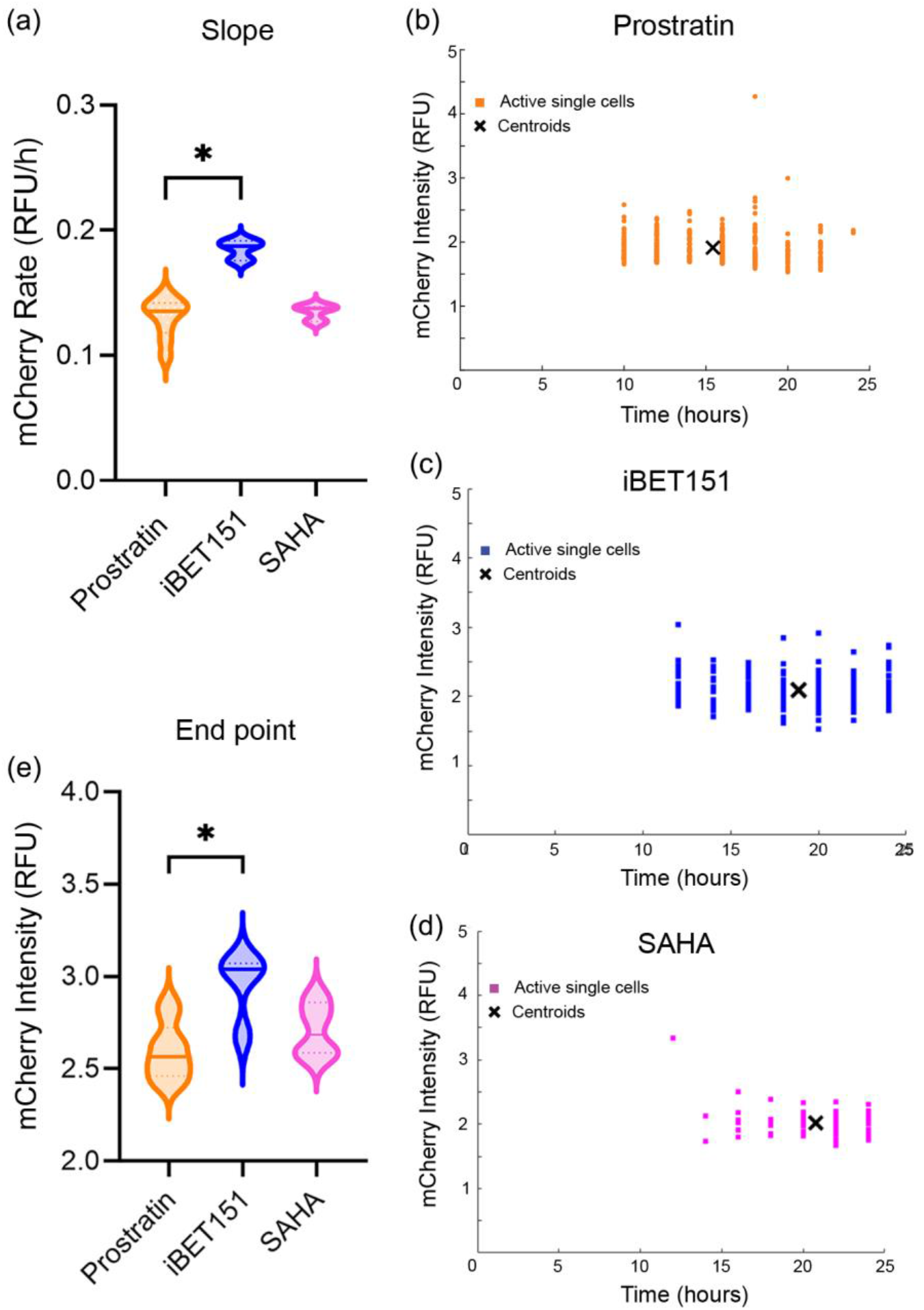
Response of single cells to LRAs. (a) Violin plot of the rate of mCherry fluorescence intensity increase for cells activated in response to the LRAs (prostratin: n=9 arrays; iBET15: n=3 arrays; SAHA: n=3 arrays). (b-d) mCherry fluorescence intensity (y-axis) at first observed time (fluorescence time) of elevated fluorescence for cells activated in response to prostratin (n=1 array 347 activated single cells) (b), iBET151 (n=1 array, 64 activated single cells) (c), or SAHA (d) (n=1 array, 46 activated single cells) for one representative experiment. Each data point represents a single cell and the crosshatch the centroid of the data points. (e) Violin plot of the fluorescence intensity at 24 after exposure to the LRAs. For panels (a) and (e), the solid horizontal line marks the median while the dashed horizontal lines the quartiles.

### 3.3 Identification of fast and slow reactivating cells after exposure to prostratin

Prostratin-activated cells were examined in greater depth by clustering the cells into groups based on the timing and intensity of mCherry fluorescence increase. Three groups were identified: non-activated (Figure 4b), “fast” reactivation (Figure 4c) and “slow” reactivation (Figure 4d). The non-reactivated group was similar to that described previously and was characterized by cells with no change in mCherry expression in response to prostratin during the duration of observation (24 h). The majority of cells (66.3 ± 5.8%) were in this category. The percentage of cells falling into either the fast and slow reactivation groups was not significantly different (p>0.999) (Figure 4e). The time at which cells began to demonstrate an increased fluorescence intensity was significantly different for the slow and fast cells (Figure 4f). By 10 h, the fast cell group exhibited an average fluorescence intensity that was significantly greater than the slow group (****p<0.0001) or the non-reactivated cells (****p<0.0001). This was also reflected in the significantly longer fluorescence time of the slow relative to the fast cells (****p<0.001, Table 2). Interestingly the initial increase in fluorescence intensity was similar for both cell groups (Figure 4h). Although over the course of reactivation, the fluorescence intensity increased at a faster rate on average in the fast cells once these cells began to reactivate compared to the slow cells (Figure 4g, Table 2). By 22 h, the average fluorescence intensity of the slow cells begin to approach that of the fast cells, and surpassed that of the fast cells by 24 h (Figure 4i, Figure S2). None of the cells expressing mCherry were observed to undergo a decrease in fluorescence during the 24 h assay, likely due to the stability of the mCherry construct (t_1/2_~ 24 h)^25^. Taken together these data suggest that the fast and slow cell groups take somewhat different reactivation pathways.

**TABLE 2:** Characteristic of fast and slow cells activated by prostratin.

| Parameter | Prostratin |  |
| --- | --- | --- |
|  | Fast | Slow |
| **Fluorescence time (h)** |  |  |
| Mean of all cells in the category | 11.2 ± 4.7 | 19.7 ± 4.4 |
| Range (longest and shortest for each category) | 4-16 | 14-24 |
| **Rate of mCherry fluorescence increase (RFU/h)** |  |  |
| Mean of all cells in the category | 0.24 ± 0.04 | 0.16 ± 0.02 |
| Range (slowest and fastest in a single cell) | 0.16 – 0.30 | 0.13 – 0.17 |
| **Cell reactivation rate (# of cells/h)** |  |  |
| Mean of all cells in the category | 14.6 ± 5.0 | 15.7 ± 5.2 |
| Range | 8.6 – 23.8 | 7.1 – 23.1 |

**FIGURE 4:**
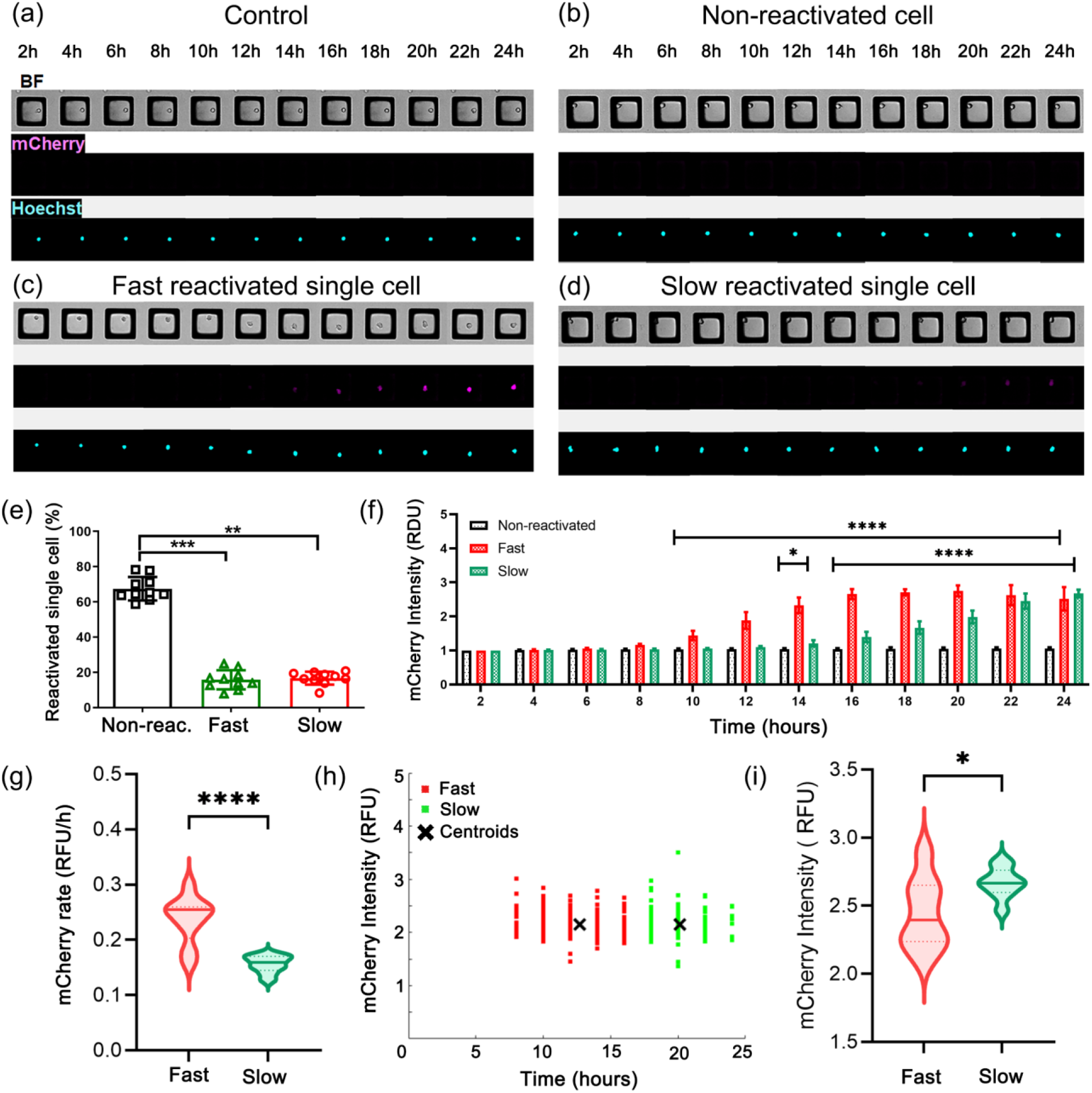
Slow and fast reactivated cells in response to prostratin. (a-d) Bright field and fluorescence images (mCherry and Hochest 33342) of a control single cell (no LRA, a), and single cells at different times after exposure to prostratin: non-reactivated (b), fast reactivated (c), and slow reactivated (d) cell. (e) Shown is the percentage of cells in the non-reactivation (gray, n=9 arrays, 5767 single cells),fast (red, n=9 arrays, 1355 single cells), and slow (green, n=9 arrays, 1355 single cells) reactivation categories. (f) Average mCherry intensity for each reactivation category over time. (g) Violin plot of the rate of mCherry fluorescence intensity increase for cells activated in response to the LRAs. (h) mCherry fluorescence intensity (y-axis) at first observed time (fluorescence time) of elevated fluorescence for cells activated in response to prostratin. Each data point represents a single cell and the crosshatch the centroid of the data points (n=1 array, 209 fast single cells and 138 slow single cells). (i) Violin plot of the fluorescence intensity at 24 after exposure to the LRAs. For panels (g) and (i), the solid horizontal line marks the median while the dashed horizontal lines marks the quartiles.

### 3.4 s.c.RNA sequencing data

Since prostratin induced a higher fraction of reactivated cells compared to SAHA or iBET151, further sorting and collection was carried out for single-cell RNA sequencing. Cells that followed a fast or slow reactivation during prostratin exposure were collected using an up-right mechanical system with a magnetic wand. Non-reactivated and control cells (no LRA) were also collected. For scRNA-seq analysis, we first performed a quality control checking the number of reads (top), the percentage of mitochondrial transcripts (center), and the number of genes being expressed (bottom), and no significant differences between groups were found (Figure 5a). Our scRNAseq data suggests that the reactivation kinetics of HIV-infected cells (fast vs. slow) is largely uncorrelated with the transcriptome of the individual cell. However, since several hours elapsed between reactivation and extraction for some cells, the transcriptomic state of the cells at the time of reactivation could have changed over time. As such, profiling cells at the time of reactivation may reveal a distinct reactivation-associated phenotype (Figure 5b).

**FIGURE 5:**
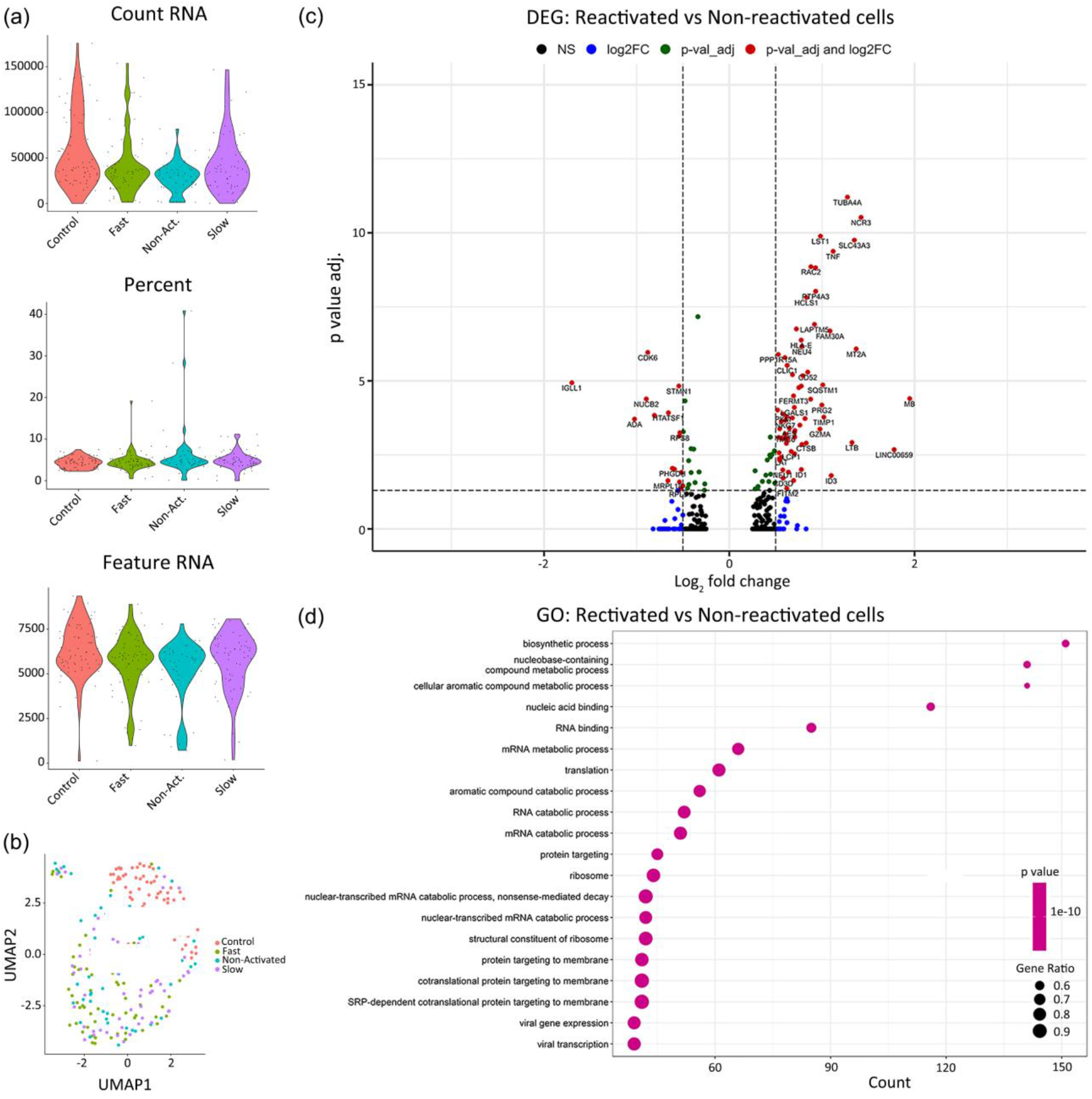
(a) scRNAseq quality control for all single cells in each group, control, non-reactivated, fast, and slow reactivating cells. (b) UMAP clusters for control (red), non-reactivated (blue), fast (green) and slow (purple) activated cells. (c) Volcano plot of significantly differentially expressed genes in reactivated cells related to non-reactivated cells. Red dots represent genes with higher expression levels with adjusted log10 p-value (0.05) and log2 fold change (0.5). The right red dots are related to upregulated genes and the left red dots to downregulated genes in reactivated cells. Green dots represent significantly different genes with an adjusted log p value < 0.05 and lower log2 fold change value. Blue dots represent genes with a greater log2 fold change but an adjusted log p-value higher than 0.05. Black dots are non-significant genes. (d) Gene Ontology (GO) analysis showing different processes of upregulated genes in activated cells compared with non-reactivated cells.

Interestingly, several differentially expressed genes (DEGs) showed higher expression in the reactivated cells than in the non-reactivated cells (Figure 5c and Figure 6). Gene Ontology (GO) analysis showed that upregulated genes in reactivated cells were related to nucleic acid binding and RNA binding (Figure 5d and Figure 7). Moreover, our data showed that catabolism and metabolism pathways are associated with viral reactivation in latently infected cells (Figure 5d and Figure 7). Immunometabolism is increasingly recognized as playing an important role in HIV expression, and this could represent a promising way to modulate HIV infection and latency.^38^

**FIGURE 6:**
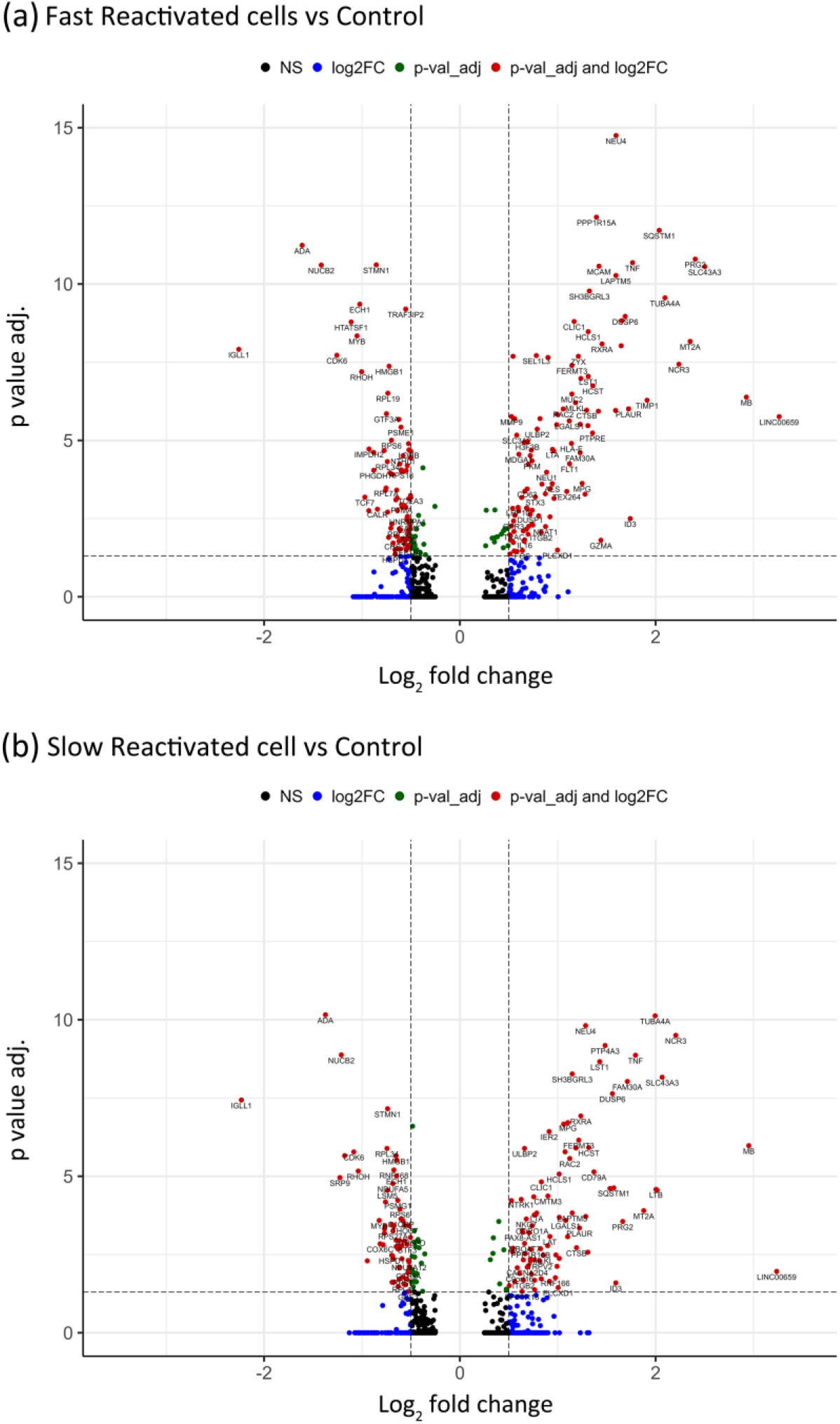
Volcano plots of significantly differentially expressed genes in (a) fast and (b) slow reactivating cells compared with control cells (no LRA). Red dots represent genes with higher expression levels with adjusted log10 p-value (0.05) and log2 fold change (0.5). The right red dots are related to upregulated genes in fast and slow reactivating cells, while the left red dots are downregulated genes. Green dots represent significantly differentially expressed genes with an adjusted log p value < 0.05 with a lower log2FC value. Blue dots represent genes with a greater log2FC but an adjusted log p-value higher than 0.05. Black dots are non-significant genes.

**FIGURE 7:**
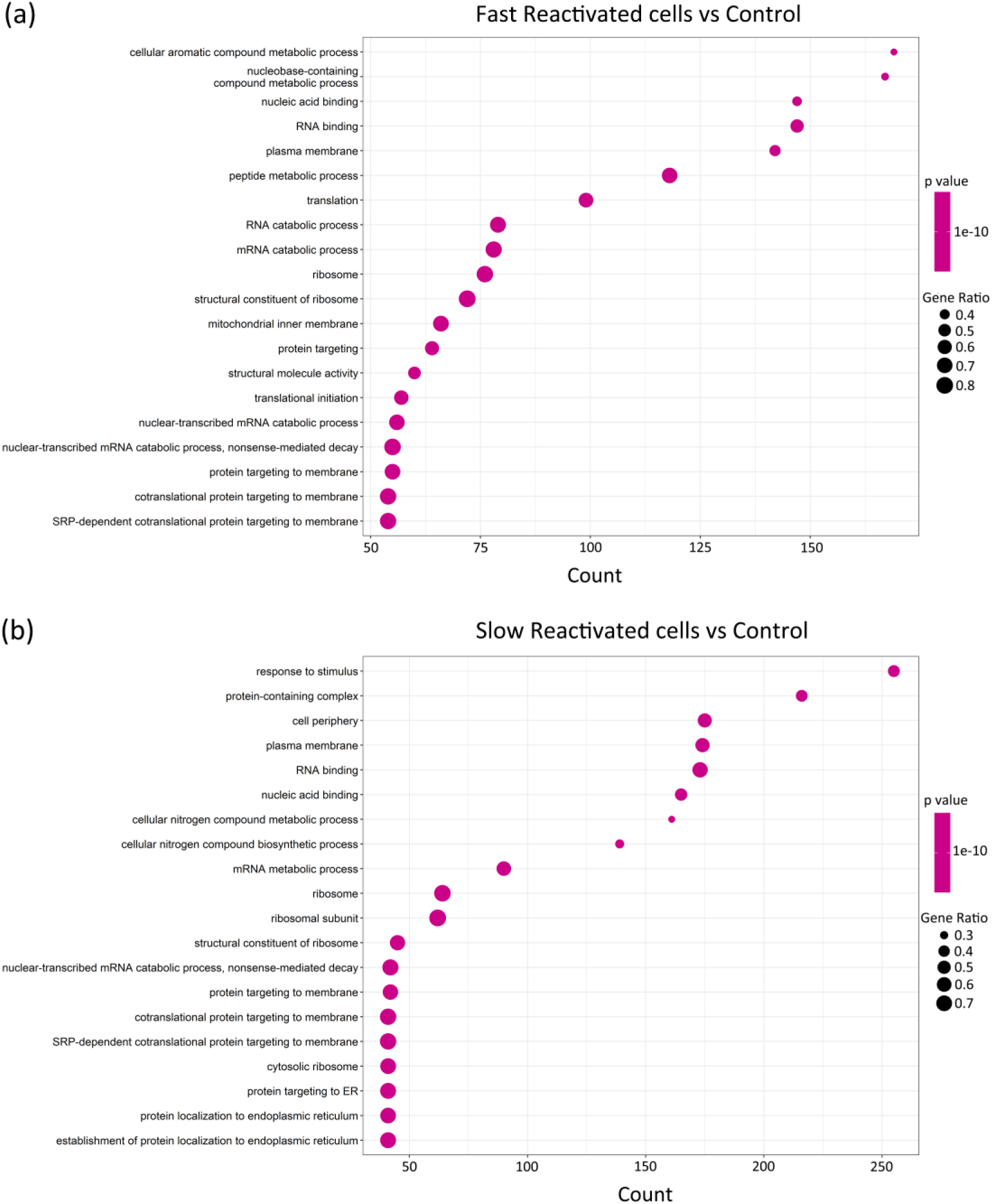
Gene Ontology (GO) analysis showing different processes of upregulated genes in (a) fast and (b) slow cells compared with control cells.

Interestingly, some of the DEGs identified by the scRNAseq analysis have known or suspected roles in HIV infection. For example, metallothionein (MT)2A was upregulated in reactivated cells (Figure cand Figure 6). MT genes are related to the immune response or defense response.^39^ The lysosomal-associated transmembrane protein 5 (LAPTM5) was also highly expressed in reactivated cells (Figure 5c and Figure 6). This gene has been implicated in the activity of the viral protein R (Vpr), which is needed for high viral load and disease progression. Vpr counteracts restriction by LAPTM5 enhance HIV infection in dendritic cells and macrophages. ^40,41^ LAPTM5 is downregulated in cell lines such Jurkat cells and primary CD4 T cells by Vpr, and this activity could be necessary for HIV replication.

Moreover, upregulated genes related to inflammation, proliferation, and immune regulation such as Linker for activation of T Cells 2 (LAT2), Tumor Necrosis Factor (TNF), and Retinoid X Receptor Alpha (RXRA) were highly expressed in reactivated cells (Figure 5c and Figure 6). T cell activation and proliferation is associated with inflammation.^42^ and HIV immune activation and inflammation increases TNF, which is involved in the regulation of cellular proliferation, inflammation, and immune regulation.^43–45^ TNF alpha also potently induces activity of NFkB, a known regulator of HIV transcription.

Downregulated genes in reactivated cells included TCF7, a transcription factor associated with quiescent memory T cells. Previous work has shown that TCF7 expression is enriched in HIV-infected CD4 T cells with low levels of viral transcription^12^. Furthermore, HTATSF1, a known regulator of the HIV transcription factor Tat was also downregulated in reactivated cells. This protein has been shown to bind the HIV genome and facilitate selective transport of HIV RNAs.^46^ The transcriptional regulator c-Myb was also enriched in this cluster, and this gene has been shown to influence HIV gene expression.^47^ Additional work to selectively target these genes in this model system may reveal a functional role in HIV reactivation from latency.

## Conclusions

Single cell approaches to study the heterogeneous responses of latent HIV reservoirs cells are critical to enable a cure for HIV. We developed microraft arrays to enable the temporal tracking of reactivating single cells followed by cell sorting and gene expression analysis. Importantly the gene expression profiles could be mapped directly to the reactivation kinetics for each cell. Cells stimulated with prostratin displayed the shortest time to reactivate HIV expression and the greatest percentage of cells reactivated compared to iBET151 or SAHA. However, cells reactivated by SAHA or prostratin displayed a slower rate of increase in mCherry fluorescence compared to iBET151. For prostratin exposed cells, sufficiently large numbers of cells could be tracked to enable clustering of the cells into non-activated, fast reactivation and slow reactivation based on their fluorescence intensity over time. The fast reactivating cells demonstrated a greater rate of mCherry fluorescence increase once the cells initiated reactivation, but reached an overall lower fluorescence intensity compared to the slow reactivating cells. Given the rapid maturation time of mCherry (t_1/2_~ 40 min) relative to the timescales of these experiments we believe that the kinetics of reactivation are accurately depicted. Given the high stability or long degradation time (t_1/2_~ 24 h) for mCherry, the kinetics with which LChiT3.2 cells return to a latent state after application of LRAs cannot be accurately observed in these experiments and will require a destabilized fluorescent protein with a fast degradation time. Lastly, our approach identified genes associated with inflammation, immune activation, and cellular and viral transcription factors. This approach and identified genes associated with latency reversal may enable selective targeting these genes in model systems to further our mechanistic understanding of HIV latency and viral reservoir towards a cure for HIV.

## Conflict of interest

NLA discloses a financial interest in Cell Microsystems, Inc. All other authors declare no conflicts.

## Funding

This work was supported by NIH grants: GM123542 (to DMM) and CA224763 (to NLA.) NIAID R01 AI143381 (EPB), NIDA R61 DA047023 (EPB).

## Author Contributions

BCL performed microfabrication and time-lapse imaging, sorting and single cell collection experiments. BCL conducted the automated clustering and kinetics analysis. AV conducted the library preparation experiments. VJ performed the s.c.RNA-sequencing analysis. BCL, NLA, conducted the discussion of the single-cell heterogeneity during reactivation. BCL, EB and DM conducted the discussion of the s.c.RNA-sequencing data. All authors revised the manuscript.

