## Supplementary Information for "Automated microarray for single-cell sorting and collection of lymphocytes following HIV reactivation"

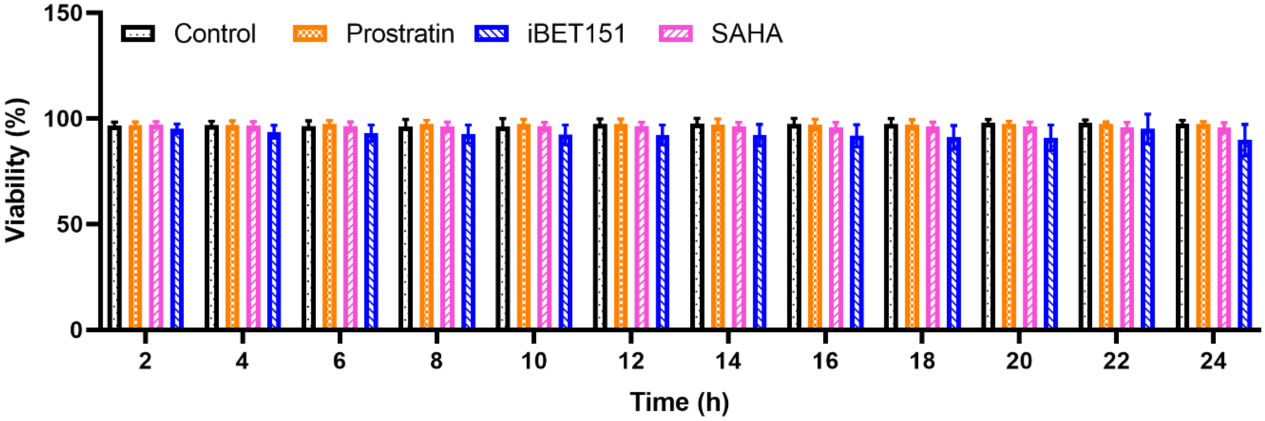


**FIGURE S1:** Lymphocytes viability after 24 hours without LRA (control, black) and under 1 µM of prostratin (orange), iBET151 (blue), and SAHA (magenta). LRA: prostratin ($p>0.995$, 10 arrays, 8768 single cells), SAHA ($p>0.848$, 4 arrays, 6354 cells), and iBET151 ($p>0.139$, 4 arrays, 5118 cells).

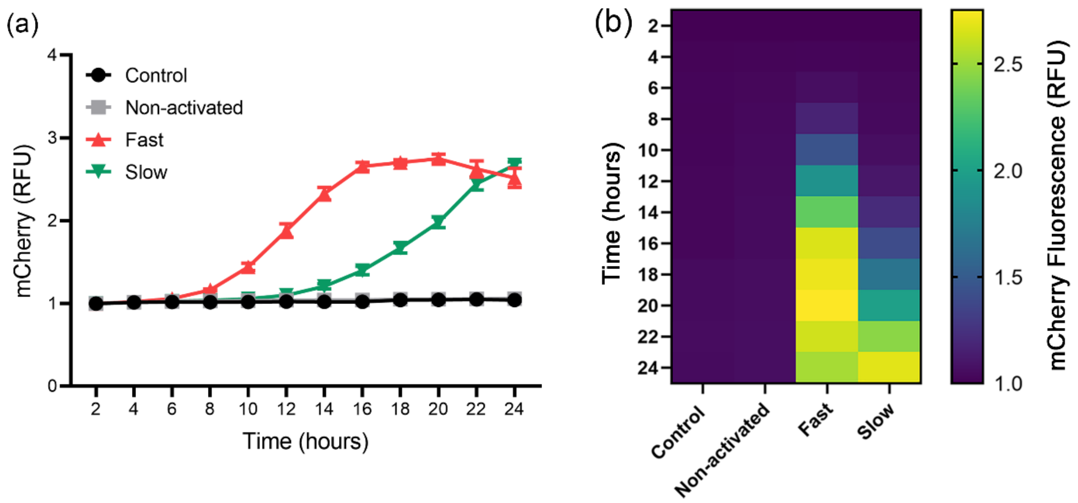


**FIGURE S2:** Variability and clustering of the mCherry intensity of reactivated HIV latency cells under prostratin. (a) mCherry intensity of each cluster over time. (b) Heat map plots showing the mean intensity values of all control, non-activated, fast, and slow cells.
